# Demography and behaviour of polygyne nests of the supercolonial ant *Cataglyphis niger*: Does kinship matter?

**DOI:** 10.1101/392522

**Authors:** Tali Reiner Brodetzki, Guy Brodetzki, Ofer Feinerman, Abraham Hefetz

**Affiliations:** School of Zoology, George S. Wise Faculty of Life Sciences, Tel Aviv University, Tel Aviv, 6997801, Israel.; Department of Physics of Complex Systems, Weizmann Institute of Science, Rehovot, 7610001, Israel.

**Keywords:** nepotism, social structure, supercolony, QR tracking, kin selection, polygyne

## Abstract

The basic ant colony is presumed to have evolved through kin selection. However, ants show a remarkable diversity in their social organization, from a monogynous-monandrous queen to the more derived states of polygyny with polyandrous queens. The existence of polygyny is an evolutionary enigma, since kin selection theory predicts that while queens should strive for reproductive monopoly, workers are predicted to favor their own matriline in rearing gynes. Using a barcoding system that enables tracking of individual interactions, along with polymorphic DNA microsatellite markers that indicate the matriline and patriline of all individuals, we demonstrate the complex social interactions in polygyne nests of *Cataglyphis niger*. *C. niger* is not only polygyne but also constitutes a supercolony at the study site. Our pioneering findings that both queens and workers are not necessarily related to each other support the supercolony structure of the population. Also in line with supercoloniality, we demonstrate that the workers contribute equally to the nest production and rearing of the queens. Unlike invasive supercolonial species, *C. niger* is native to Israel, raising questions about the driving forces, apart from kin selection, that stabilize this society.

## Introduction

In the Middle Ages, the chastity imposed upon Catholic popes led to an inclusive fitness behaviour of appointing their nephews as cardinals, who would later succeed them as popes. This behaviour is known as nepotism. Similarly, monogynous ant colonies, the postulated basic colony structure, presumably evolved via kin selection (Hamilton, 1964; Hölldobler and Wilson, 1990; Hughes *et al*., 2008; Boomsma, 2009). Notwithstanding, more derived states in which the colony is composed of multiple queens (polygyny) that may also be multiply inseminated (polyandry), are very common and present a conflict for the inclusive fitness theory. Polygyny poses two problems: from the queens’ perspective, queens are predicted to compete for monopoly in reproduction; while from the workers’ perspective, workers are predicted to favor their own matriline in rearing gynes (young unmated queens; Keller, 1993). The social consequences of polygyny in ant colonies vary, ranging from an equal reproductive share among queens to functional monogyny (i.e., only one queen reproduces while all other queens behave like workers); and from complete openness to incoming queens to a limited acceptance of only related queens (Hölldobler and Wilson, 1977). Species may exhibit social polymorphism with this respect, which is mostly believed to be the result of multiple selective pressures such as ecological constraints on nesting success, predation, and increased genetic diversity. Social polymorphism is also usually associated with profound changes in life-history strategies and dispersal behaviour (Bourke and Franks, 1995; Keller, 1995; Pamilo and Crozier, 1996; Ross, 2001; Chapuisat, Bocherens and Rosset, 2004). Inclusive fitness theory also assumes favorable discrimination of closely-related individuals (e.g., nepotism), unless between-matriline competition hampers total colony reproductive output. This discrimination is probably mediated through kin recognition cues, similar to nestmate recognition cues. Although, recognition cues exist in ants (Soroker *et al*., 1994; Lahav et al., 1998) they seem to play a role in within-colony kin discrimination only in rare cases (El-Showk *et al*., 2010). Additionally, in most cases relatedness in these polygyne nests is low thus raising the question of the benefits of the inclusive fitness theory. Indeed, to date the occurrence of nepotism in eusocial insects, specifically in ants, has mostly been rejected (DeHeer and Ross, 1997; Holzer et al., 2006; Zinck, Châline and Jaisson, 2009; Friend and Bourke, 2012), except for *Formica fusca* (Hannonen and Sundström, 2003) but see the criticism by (Holzer *et al*., 2006; e.g. selective brood mortality rather than preferential brood rearing), and *Leptothorax acervorum* ants (Gill and Hammond, 2011). The extent of nepotism in eusocial insects is thus still unclear and there is a need for actual behavioural assays. Boomsma and d’Ettorre (2013) suggested that within-colony kin discrimination might be maintained in a secondary polygyny where there is a high queen turnover and fluctuating relatedness in the colony.

The ant genus *Cataglyphis* comprises over one hundred described species that are distributed across the Palaearctic region in arid environments (Agosti, 1990). *Cataglyphis* exhibits a high social polymorphism, from strict monogyny (*C. sabulosa, C.bicolor, C. emmae and C. hispanica*) through facultative polygyny (*C. velox and C.livida*) to polygyny (*C. niger and C. mauritanica;* (Lenoir *et al*., 2009; Boulay *et al*., 2017). The genus also demonstrates a diversity of reproduction modes (sexual and asexual), which enables us to investigate the evolution of sociality. It is therefore highly important to understand the mode of social organization for each species. Specifically in polygyne nests, it is important to understand the within-nest interactions since there are multiple possibilities of nest foundation and relatedness asymmetries (Keller and Reeve, 1995).

Here, we focus on a population of *C. niger* that constructs polygyne nests organized as a supercolony (Leniaud *et al*., 2011). In such population structure, brood and workers are often translocated to other nests in the population according to the changing needs of the widespread colony. This complex social system raises multiple thoughts about the mechanisms leading to this evolutionary stable status. Nepotism was investigated and rejected in the imported fire ant, *Solenopsis invicta*, which has a similar population and social structure in its invasive range (DeHeer and Ross, 1997). However, *C. niger* is a native species, and therefore could behave differently and possibly demonstrate within-colony kin discrimination more than an invasive species.

Using a barcode system for tracking individual movement, we monitored worker-worker and workers-queen interactions in polygyne nests. We also genotyped the workers and queens in order to assess the within-nest relatedness. For the first time, to the best of our knowledge, we have obtained extensive data on both individual interactions and relatedness between the tested individuals, revealing the complex relationships that exist in polygyne nests, and rejecting the possibility of nepotism.

## Methods

### Ant collection and maintenance

Three complete polygyne nests (nests 677, 868, and 869 possessing 3, 3 and 4 queens, respectively) of *C. niger* (*niger* haplotype, as described in Eyer 2017) were excavated from the Tel-Baruch population in the semi-stabilized sand dunes of Tel Aviv, Israel. Prior to any manipulation, nests were kept in the lab for about a month under constant temperature and humidity and provided with water and a diet according to Bhatkar and Whitcomb (1970), supplemented with tuna fish. For observations, a subset of randomly selected workers (n=200 for each of the 3 nests) and the queens were housed in artificial nests (20cm x 30cm) closed with an IR filtered roof and provided with water and food *ad libitum*. Workers and queens were each marked with individual 2mm QR barcodes using skin adhesive (SAUER-HAUTKLEBER, Manfred sauer GMBH). Using a Canon T3i we took snapshots of the nest every 2 seconds, for 10,000 continuous frames (approximately 5.5 hours). The following observation protocol was performed for each of the three nests. Three days after tagging the first session began, constituting free roaming queens and workers, and lasted for 3 days. In the following session the queens were tethered at the petiole with fine nylon string (2-3 cm long), restricting each to a limited space in a different corner of the nest, while allowing free movement of the workers throughout the nest. At the end of this session the queens were released for three days, after which a second session of tethered queens began, in which the queens were placed at different corners from the previous session.

### Statistical analysis

Behavioural analyses were performed using BugTag, a barcode analysis system (developed by *robiotec*). The tags enabled continuos monitoring of the location of each individual ant. The direction of the tags was aligned to match the position of the ant’s head, indicative of the direction at which it faced. Workers presenting low detection rates (their identity was certain in less than 30% of the 10,000 frames) were excluded from the analyses. In cases where missed detections occurred in segments that were either fewer than five frames (less than 10 second apart), or if the change in the individual’s position before and after the detection gap was less than 2cm in distance, the position was interpolated using linear regression. Thus, the numbers of workers in each analysis (2 replicates) were 137 and 122 for nest 677; 149 and 161 for nest 868; and 161 and 159 for nest 869.

Using MATLAB, we defined a circular virtual arena (2cm radius) around each queen, hence termed “retinue space”. Workers were considered to engaged in retinue around the queen if they were positioned in the “retinue space” while facing the queen for over 10% of the observation time. Ants that only occasionally interacted with a queen or sporadically shared her roaming space were not considered as involved in retinue behaviour. The constancy of these groups over time, in terms of ant identity, and the fidelity of each worker to a specific queen or task were also measured. A Permutation analysis was used to test the significance of the ants’ positions between the different queens’ spaces. During the manipulation, the queens were shuffled in the nest (tied queen iteration 1 & 2) and the ants reoccupied retinue positions. Computationally, we define a score for each possible one-to-one assignment (Heyman *et al*., 2017), that score takes in to account the changes in worker position between iteration 1&2. This measure varies between 0 and 1. If it occurs that each ant returned to exactly the same queen’s space after the manipulation (including fraction of time spent) then the actual experimental assignment will be scored as 1. If ants’ retinue other queens or do not retinue at all the score can be expected to be closer to the minimal value of 0. Thus, this score is given both to the actual ant assignment as projected by the experiment and the random permutations of their locations. If the actual score is different than the mean permutation score than the ants are choosing their positions; either to stay with the same queen or rather choosing to retinue another queen. However if the real score is no different than the permutation score than ants’ retinue is random.

### Genetic analysis

At the end of the experiment, all the queens and workers were sacrificed for genetic analyses. For nest 868 we analyzed the 3 queens and all the 165 workers in the experimental nest. For the remaining two nests (677and 869) we analyzed the queens and a subset of workers (18 and 28, respectively), which included the top three “retinue” workers for each queen (the minimal number of retinue workers for all of the queens in all of the sessions) along with randomly selected nestmates. Queens’ ovaries were dissected to assess their degree of activation. Under a binocular we compared the number of reproductively active ovarioles and the number and size of oocytes in them, creating an activation index as follows: **0** - non-activated ovarioles; **1** - 2 to 4 activated ovarioles with slightly developed oocytes; **2** - 4 activated ovarioles with medium oocytes; **3** - all ovarioles with large, ready to lay eggs). We also kept the spermathecal content of each queen for genetic analyses in Ringer’s buffer at -80°c until DNA extraction. DNA was extracted with 5% CHELEX (BIO-RAD) and amplified with 7 microsatellite markers that had been previously designed for *C. hispanica* (Ch23; Darras, Kuhn and Aron, 2014), *C. cursor* (Cc51,Cc89, and Cc99;Pearcy *et al*., 2004), and *C. niger* (Cn02, Cn04, and Cn08; Saar *et al*., 2014) using Type-it PCR mix (QIAGEN). PCR products were sequenced by ABI3500 genetic analyzer and genotype analyses were performed with GeneMarker. Relatedness was evaluated with Coancestry (Wang, 2011) and ML Relate (Kalinowski, Wagner and Taper, 2006). We checked for possible correlation of relatedness to the mean distances between all the individuals in the nest (workers and queens) using SOCPROG 2.8. Network analysis was done using UCI6 and NetDraw.

## Results

### Nests compositions and within-nest relatedness

Each experimental nest had 3-4 queens (667-3; 868-3; 869-4), all of which were multiply inseminated by 3-7 males. All the queens in the experiment (n=10) had well developed ovaries with ready-to-lay eggs (level 3), except for one queen in nest 868 (**Q**ueen 3- tag # 552) and one queen in nest 869 (Q4- tag # 1189) which had medium-size oocytes (level 2). Table 1 presents the level of relatedness among the queens in the experimental nests. In nest 869 three of the four queens were highly related (Relatedness (R)>0.5; full sisters or daughters), while the 4^th^ queen was completely unrelated (R<0; not different from relatedness=0). In nest 677 two of the queens were slightly related (0.2181; half sisters or cousins), while the 3^rd^ was not related to either of the other two (R<0.1). All three queens in nest 868 were completely unrelated (R<0.1).

**Table 1:** Relatedness values between queens

| Nest - Queen | 869 Q2 | 869 Q3 | 869 Q4 |
| --- | --- | --- | --- |
| 869 Q1 | 0.5304 | - 0.1901 | 0.5362 |
| 869 Q2 | -- | 0.0626 | 0.6122 |
| 869 Q3 | -- | -- | - 0.0165 |
|  | 667 Q2 | 667 Q3 | -- |
| 667 Q1 | 0.2181 | 0.047 | -- |
| 667 Q2 | -- | 0.0794 | -- |
|  | 868 Q2 | 868 Q3 | -- |
| 868 Q1 | - 0.1573 | 0.0873 | -- |
| 868 Q2 | -- | - 0.3619 | -- |

To assess nest demography, we genotyped all the workers and queens from nest 868 that had participated in the tagging experiment (three queens and 165 workers). Since these workers were randomly selected out of the approximately 1,000 workers that populated the nest, we assume that they reflect a good assessment of the genotype distribution within the entire nest. The genotype analysis revealed that 9.6% of the workers were highly related to each other (number of relationships/total possible relationships of Relatedness value>=0.5), 14.8% were moderately related (0.25<=R<0.5), 19.3% had low relatedness (0.125<=R<0.25), and 56.3% were unrelated to any of the other workers in the nest (R<0.125). A network analysis of relatedness values with ties indicating relatedness greater than 0.25 and nodes sized according to centrality measures is presented in supplementary Figure S1. Relatedness distribution of the workers to the different queens was as follows: Highly related workers (R >=0.5, possible daughters or full sisters) **Queen** 1=10.3%, Q2=9.1% (of which 1 worker was similarly related to both Q1 and Q2), and Q3= 6%. Moderately related workers (0.25<=R<0.5, possible sisters) to the different queens was Q1=14.5%, Q2=15.8% (of which 4 workers were similarly related to both Q1 and Q2), and Q3= 2.4%. The remaining 63 workers were not related to any of the three queens, (R< 0.25). Overall, nest relatedness for nest 868 was 0.058, and average relatedness of the workers to each of the queens was similar (0.089±0.004). These results are slightly lower than the within-nest relatedness of 0.112+0.009 of the entire population (based on 129 workers from 6 nests; data not presented).

For nests 677 and 869 we genotyped a subset of workers as presented in Table 2. In nest 677, 27.7% of the 18 selected workers were highly related to at least one of the queens (relatedness >=0.5): Q1=5.5%, Q2= 22.2%, and Q3=0; and 38.9% of the workers were moderately related (0.25<=Rx<0.5, possible sisters): Q1=27.7%, Q2=5.5% (one worker who was equally related to Q1), and Q3= 11.1%. The remaining 38.8% of the workers were not related to any of the queens (R<0.25). In nest 869, 28.6% of the 28 selected workers were highly related to at least one of the queens (R >=0.5): Q1=7.1%, Q2=3.6% (one worker who was similarly related to Q1 and Q4), Q3=7.1% (one worker was similarly related to Q4), Q4=17.9%. Moderately related workers to queens were as follows: Q1= 14.3% (one worker similar to Q4), Q2= 21.4% (one worker similar to Q3 and one to Q4), Q3= 7.1%, Q4=17.9%. The remaining 42.9% of the workers were not related to any of the queens.

**Table 2:** Number of workers that have relatedness values of 0.25/0.5 to any of the queens. N= 18 for nest 677 and 28 for nest 869. ^12*º^wy^ indicate individuals that are related to both queens.

| NEST_QUEEN | RELATEDNESS LEVEL |  |
| --- | --- | --- |
|  | 0.25 | 0.5 |
| 677_Q1 | 6<sup>12</sup> | 1 |
| 677_Q2 | 5<sup>2</sup> | 4 |
| 677_Q3 | 2<sup>1</sup> | 0 |
| 869_Q1 | 6<sup>12*o</sup> | 2* |
| 869_Q2 | 7<sup>1*o^wy</sup> | 1* |
| 869_Q3 | 4<sup>1^w</sup> | 2<sup>1</sup> |
| 869_Q4 | 10<sup>12wy</sup> | 5<sup>1</sup> |

### Identity and genotype of “retinue workers”

The first set of experiments was performed in an “open-range” arena where the queens and workers could roam freely. Figure 1A depicts network analyses of the mean distances between the individuals in the experimental nest 868. The network presents only the ties between individuals that were in high proximity to each other over time (3 days; proximity<mean+2SD), individuals that did not have such ties are represented at the left side of the network. The three queens as well as a small number of workers were highly central in the network: i.e., they had close interactions with many individuals. These workers can be considered as ‘retinue workers’. Queens also tended to stay very close to each other. It is also evident that, over time, many of the workers did not maintain close proximity to each other (who are presented in fig. 1A had few or no ties; their proximity to one another was below 2SD). Figure 1B depicts a network of relatedness of the ‘retinue’ workers only with the queens, demonstrating only ties higher than 0.25 (moderately to highly related). Queens were not very central with respect to relatedness values, with only a few of their daughters among the retinue workers. Furthermore, the retinue workers who were central in the proximity network were not very central in the relatedness network and not highly related to others in the nest. Aligning relatedness results and mean proximity locations of all the workers and queens revealed no clear correlation (i.e., neither workers nor queens interacted more with highly related individuals; Mantel Z test= -0.005, p=0.75).

**Figure 1:**
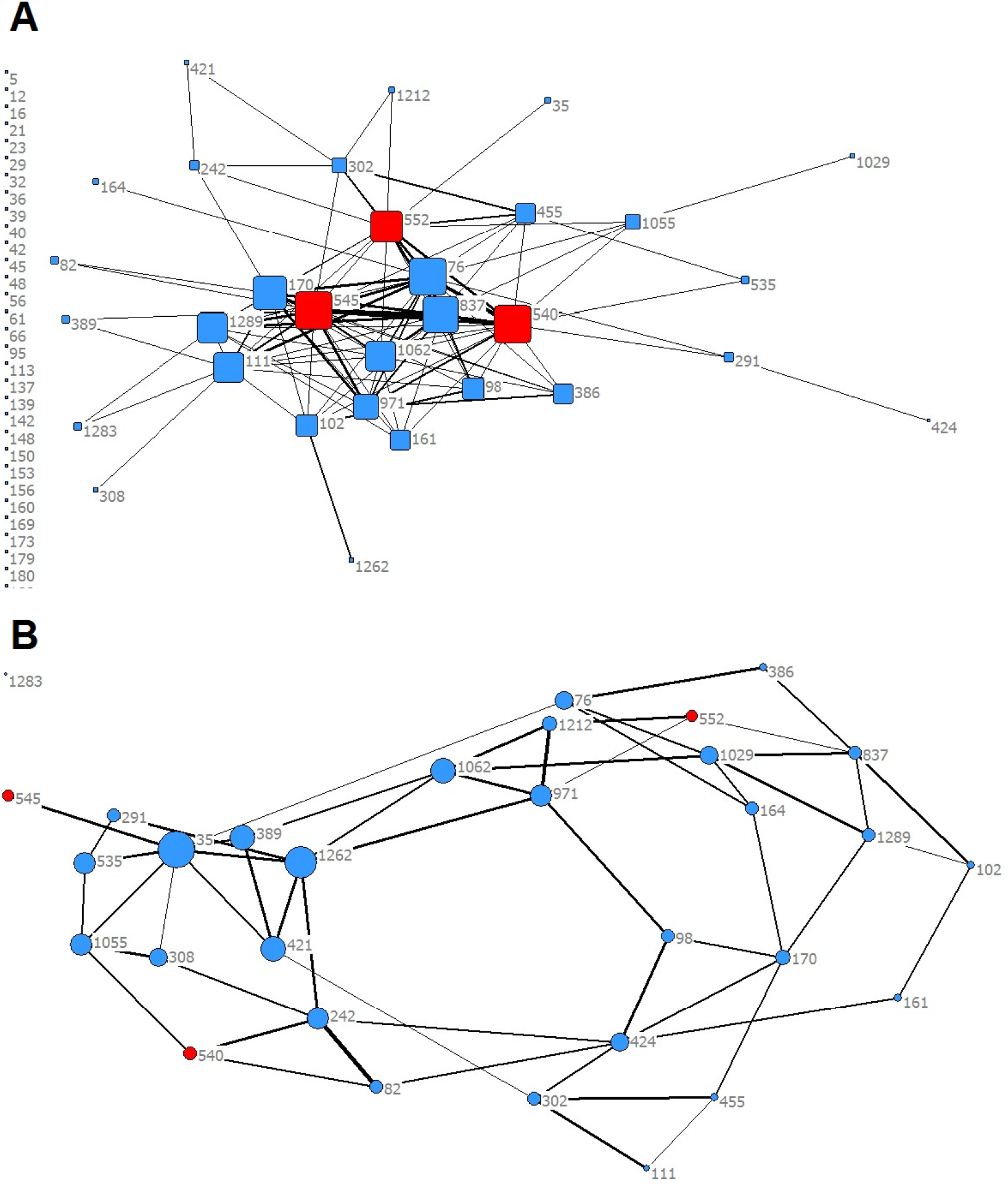
A) Network analysis of the averaged minimal distance (proximity) between the ants in nest 868 (threshold of proximity< 2*SD). Queens coloured in red. The size of the square indicates its centrality (how many relationships it contains). The thickness of the lines indicate the strength of the tie= average proximity between individuals. B) Network analysis of relatedness values of the retinue ants in nest 868 showing ties larger than 0.25). Queens coloured in red. The size of the circle indicates its centrality (how many relationships it contains). The thickness of the line indicates the strength of the tie = relatedness value between individuals.

In the “open-range” arena queens tended to be near each other, which made it difficult to determine precisely which worker was tending to which queen. Therefore, to assess whether workers preferred specific queens, we spaced out the queens within the nest by confining each to a specific location in the nest while allowing workers free movement, and thus enabling them to selectively retinue a queen. Queens’ retinue consisted of 3.86±0.15 (±SE) workers (range of averages 2.8-4.9 in workers attending a queen throughout the entire session; Fig. S2). There was no preference over time for a specific queen, although queens with smaller oocytes tended to have a smaller retinue group, this was not significantly different than the retinue groups of the other, more fecund, queens.

No workers in any nest displayed an apparent preference for a specific queen. Workers varied in the time they spent interacting with the different queens (Fig. 2A), but most did not have prolonged interactions. Approximately a quarter of the workers spent more than 10% of their time in one of the queens’ retinue space, with the maximal proportion of time spent next to a specific queen being 30%. Workers were rarely loyal to the queens they initially attended, but rather directed their retinue behaviour towards the other queens (Fig. 2B). Worker switches between queens occurred both during the same session as well as between the two iteration sessions. Worker preference for the retinue of a specific queen did not differ from the random permutations of workers’ preferences for the different queens, permutation scores were almost the same as the actual experimental scores (nest 677: 0.524-0.523; nest 868: 0.495-0.495; nest 869: 0.436-0.404; Fig.2C shows the results for nest 868).

**Figure 2:**
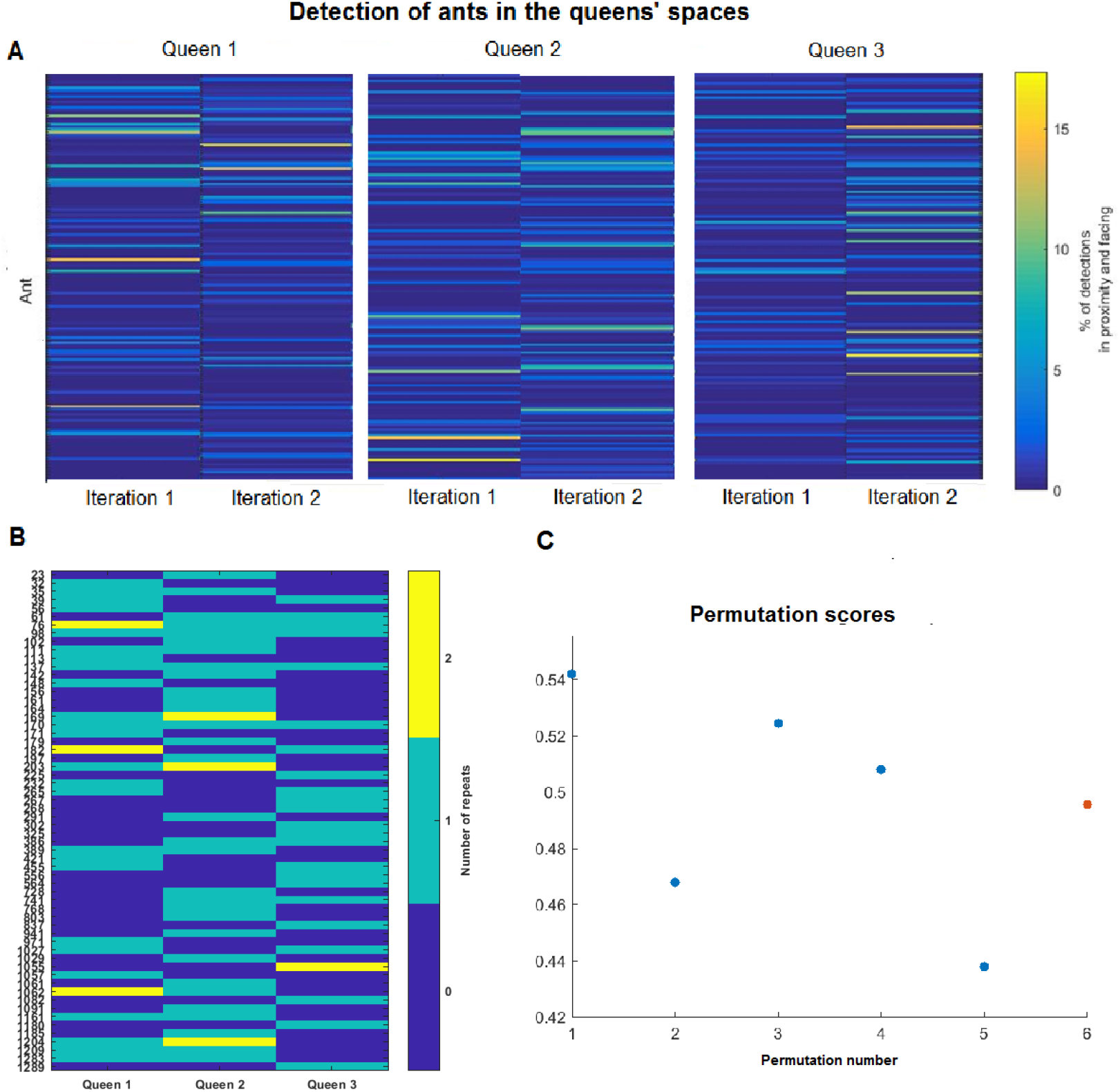
A) The proportion of time a worker interacted with a specific queen, normalized to each of the worker’s detection success in nest 868. Iterations of tethered queens were spaced by a period during which they were free. B) Workers that attended the queens for more than 10% of the time. Dark blue the workers did not attend a specific queen; light blue they attended the queen for only one repeat of the experiment; yellow they attended to the queen in both repeats. C) Permutation scores of 5 permutations for nest 868, in red the actual experimental score 0.495= mean permutations score.

The relatedness of the top three “retinue” workers (which spent more than 10% of their time in retinue) to the attended queens ranged between -0.29 to 0.54 (Fig. 3), the average of these are similar to the average relatedness of all of the workers in the nest to each of the queens, this also demonstrates that the queens’ attendants were not necessarily related to the attended queen.

**Figure 3:**
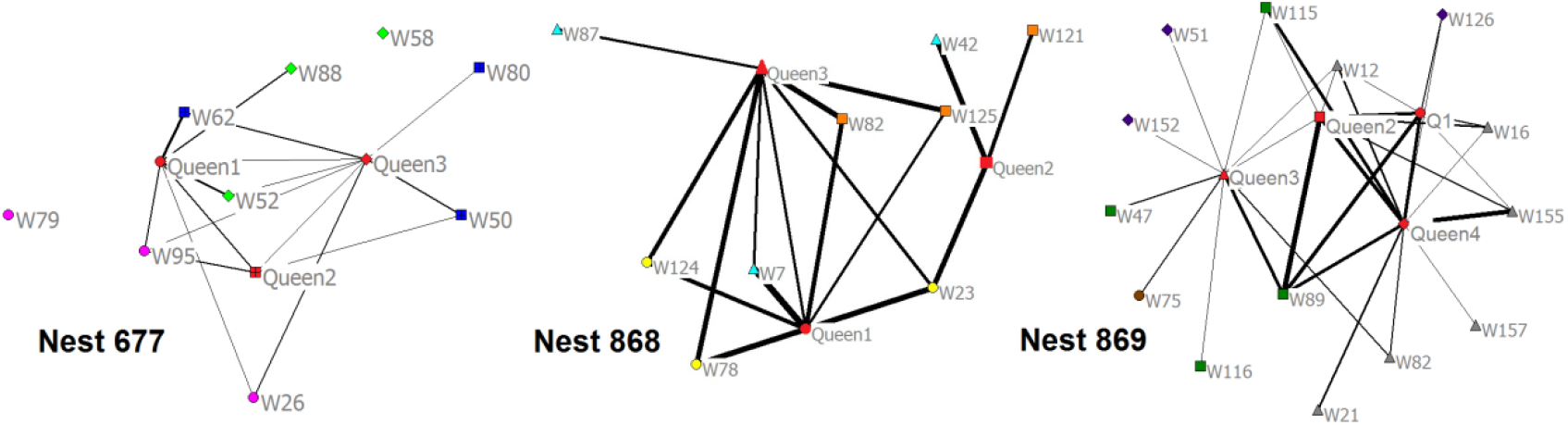
The level of relatedness between individual ants; line thickness and length represents the level of relatedness, colour and shape represent the different retinue groups of the queens, queens coloured in red (W=worker; Q= queen). Queens and their worker attendants (for each queen three of the workers that spent the most time next to the queen; relatedness under zero was emitted from the network).

## Discussion

Interactions between nestmates in an ant colony have been the focus of many recent studies, all attempting to uncover the mechanisms of ‘self-organization’, a hallmark in the evolution of social insects (Pinter-Wollman *et al*., 2011; Mersch, Crespi and Keller, 2013; Heyman *et al*., 2017). Here, for the first time, we combined individual tracking with genetic data in order to elucidate the possible effects of relatedness and kin selection on worker in-nest behaviour.

Polygyny is still an evolutionary enigma with respect to its adaptive value. Several explanations have been proposed for the mutual benefits of this social dynamic, from high relatedness of the co-inhabiting queens (the inclusive fitness explanation; Giraud *et al*., 2001) to robustness of the colony (higher genetic diversity, and colony size explanations; Keller, 1995; Hughes, Ratnieks and Oldroyd, 2008; Heinze and Rueppell, 2014). In such complex societies in which relatedness varies greatly between individuals, often reaching that of a non-eusocial non-viscous population, we would expect to see both altruistic and nepotistic interactions, like those commonly found in many vertebrate societies (Eberhard, 1975; Clutton-Brock, 2002; Hatchwell, 2010). Theoretically, nepotistic interactions should be favored unless they negatively affect the society’s overall fitness.

Additionally, in the present study, we examined whether relatedness influences the dynamics of the nest (in terms of worker behaviour) in this specific case in which the polygyne nests is also organized as a supercolony. As expected from a unicolonial system, we found that, in *C. niger* both inter- and intra-colony relatedness was very low; with only about a third of nestmate workers being related to the level of sisters or half-sisters. It was argued by Helanterä *et al*. 2009, that in order to maintain such a unicolonial structure it is necessary to have nepotism as otherwise there is a chance of the social structure disintegrating over an evolutionary time scale. However, we could not detect any indication of nepotism in our system.

Relatedness between the queens was also rather variable, ranging from sisters to un-related co-nesting queens. This suggests that this species exercises a dependent colony foundation, conforming to the usual practice in polygyne nests (Cronin *et al*.,2013). Some of the queens return to their natal nests after mating or are transferred by workers to a new nest (budding; in the case of queens that are highly related), while other queens (with low relatedness) disperse and join existing nests (this could also be explained by colony fission). From our observations at the collection site, we observed only a few newly-mated queens that were digging, presumably attempting to found a colony independently, and we also located very few nests with a single queen in them (unpublished data). Dependent colony foundation seems thus to be a preferred strategy in this population.

Our results also verify for the first time, in this specie the existence of a functional polygynous nest. Assessment of the ovarian status as well as the relatedness between queens and workers indicated that all the queens had contributed similarly to the nest population, thus establishing a true functional polygyny. Our finding that the queens were multiply inseminated further contributes to the low relatedness. Finally, we observed frequent movement between nests of workers and brood. While we cannot exclude that this was simply nest relocation, we interpret it here as free movement of workers, brood, and queens between the nests of the supercolony, according to changes in the available resources (Robinson, 2014).

The queens in our experimental nests were attended by the same number of workers on average, and also demonstrated similar oocyte development. These retinue workers comprised only a small proportion of all the workers in the nest suggesting the existence of a retinue caste, similarly to that found in honey bees (Seely, 1979), and to the nurses caste described in *Camponotus fellah* (Mersch, Crespi and Keller, 2013). The retinue workers were not closely related to the queens, and also apparently attended the different queens at random rather than being loyal to a specific queen. Indeed, worker locations were not significantly different in comparison to random permutations. A similar finding holds true for the relatedness between attendants and attendees. Nestmates did not associate more with highly-related individuals, irrespective of whether they were workers or queens. All these emphasize the existence of stochastic intra-colony interactions, both with respect to worker-queen and worker-worker interactions (i.e. the interactions do not differ from those expected at random), and thus refutes any possibility of nepotistic behaviour.

However, is nepotism such a parsimonious theory in this case? Under nepotism, we presume an unequal distribution of resources in favor of specific matrilines, which would pertrube normal colony development, rendering polygyny evolutionary unstable. On the other hand, if nepotism is maladaptive, what is the driving force for workers to keep tending non-kin, and what are the mechanisms (if any) that prevent the downgrade of non-kin-biased worker behaviour. This evolutionary conflict may explain why polygynous supercolonies are so rare, and the benefits are mostly short lived and extinguish along an evolutionary timescale. For this kind of social structure to maintain stability there should be other, ‘hard-wired’, mechanisms involved, such as the ‘social chromosome’, a non-combining chromosome that exists only in polygynous populations (Wang *et al*., 2013; Purcell *et al*., 2014). Future work should attempt to elucidate the possible mechanisms that keep these populations stable, such as worker fixed sterility.

## Acknowledgements

This project was funded by the Israel Science foundation (ISF grant # 844/13 to AH). We would like to thank Oded Schor and Yael Heyman for help with designing and analyzing the tracking methodology. We would also like to thank P.A. Eyer, Margarita Orlova, and Tovit Simon for help with sample collection and technical assistance.

## Authors’ contributions

TRB, OF and AH conceived the ideas and designed the methodology; TRB collected the data; TRB analyzed the genetic data; TRB and GB analyzed the tracking data; TRB and AH led the writing of the manuscript. All authors contributed critically to the drafts and gave final approval for publication.

## Data Accessibility

Microsatellite genotypes, CSV files of ants’ tracking and MATLAB analysis scripts will be available at: Dryad doi: to be announced after acceptance.

## Supplementary figures

**Figure S1:**
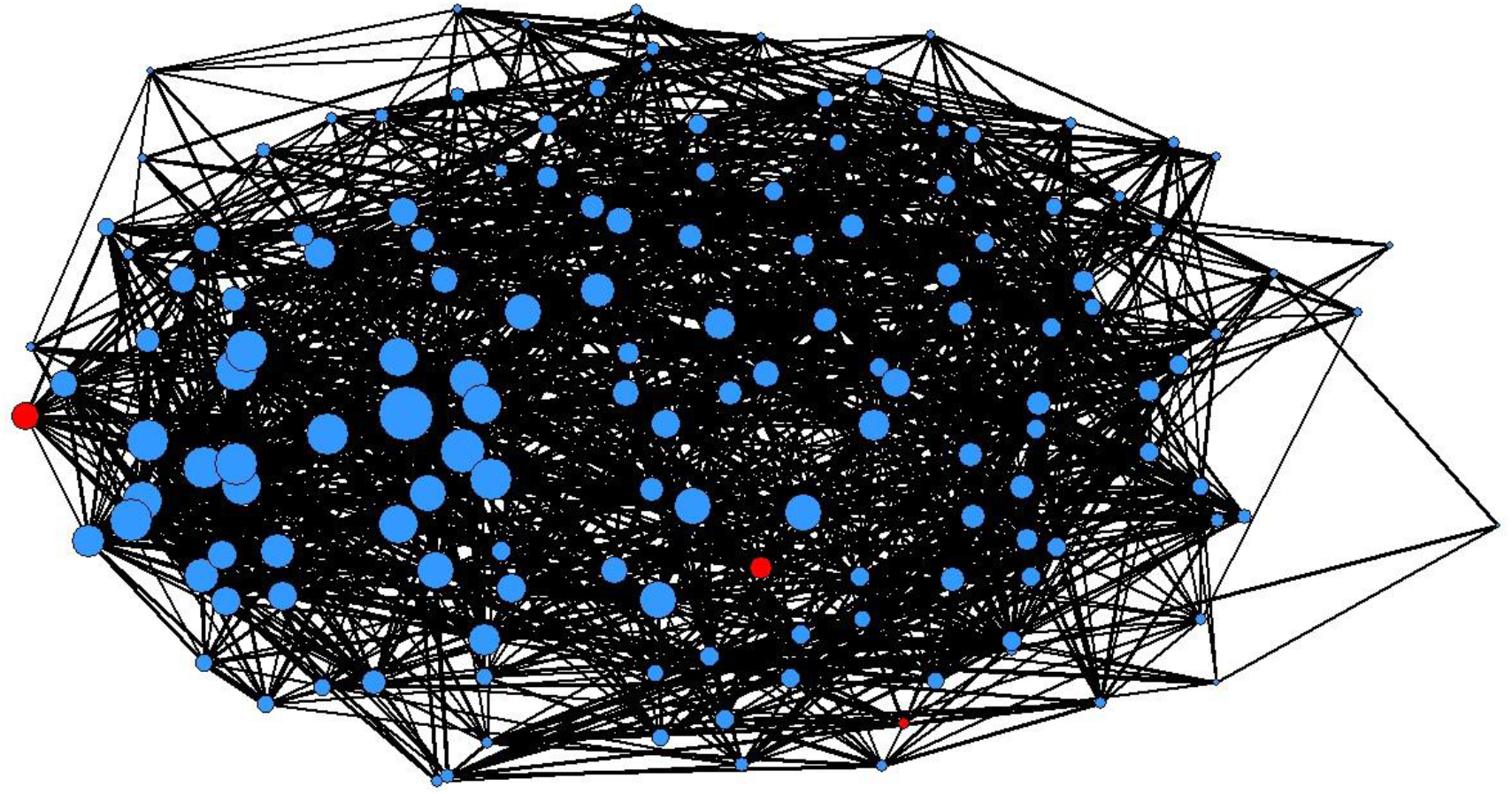
Network analysis of relatedness values of the ants in nest 868 (with a threshold of 0.25). Queens coloured in red. The size of the circle indicates its centrality (how many relationships it contains). The thickness of the line indicates the strength of the tie = relatedness value between individuals.

**Figure S2:**
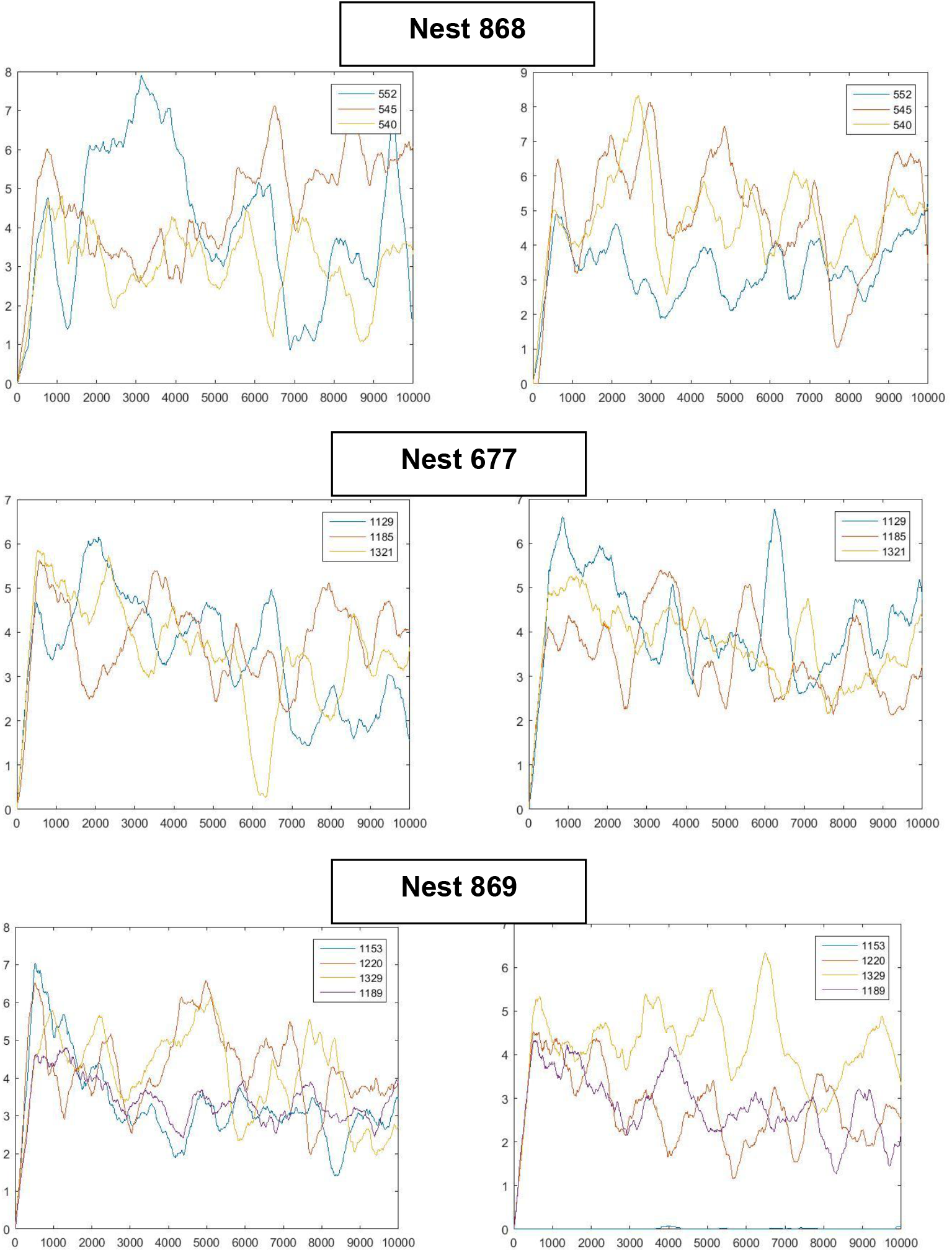
Assessment of queens’ retinue, number of workers around each queen averaged for 500 frames. Average retinue was 3.86 (SE=0.15; range:2.8-4.9) workers for each queen. In nest 868; queen 552 in blue had smaller oocytes, in nest 869; queen 1189 in purple had smaller oocytes.

